# Assembly of chloroplast genomes with long- and short-read data: a comparison of approaches using *Eucalyptus pauciflora* as a test case

**DOI:** 10.1101/320085

**Authors:** weiwen wang, Miriam Schalamun, Alejandro Morales Suarez, David Kainer, Benjamin Schwessinger, Robert Lanfear

## Abstract

**Background:** Chloroplasts are organelles that conduct photosynthesis in plant and algal cells. Chloroplast genomes code for around 130 genes, and the information they contain is widely used in agriculture and studies of evolution and ecology. Correctly assembling complete chloroplast genomes can be challenging because the chloroplast genome contains a pair of long inverted repeats (10–30 kb). The advent of long-read sequencing technologies should alleviate this problem by providing sufficient information to completely span the inverted repeat regions. Yet, long-reads tend to have higher error rates than short-reads, and relatively little is known about the best way to combine long- and short-reads to obtain the most accurate chloroplast genome assemblies. Using *Eucalyptus pauciflora*, the snow gum, as a test case, we evaluated the effect of multiple parameters, such as different coverage of long (Oxford nanopore) and short (Illumina) reads, different long-read lengths, different assembly pipelines, and different genome polishing steps, with a view to determining the most accurate and efficient approach to chloroplast genome assembly.

**Results:** Hybrid assemblies combining at least 20x coverage of both long-reads and short-reads generated a single contig spanning the entire chloroplast genome with few or no detectable errors. Short-read-only assemblies generated three contigs representing the long single copy, short single copy and inverted repeat regions of the chloroplast genome. These contigs contained few single-base errors but tended to exclude several bases at the beginning or end of each contig. Long-read-only assemblies tended to create multiple contigs with a much higher single-base error rate, even after polishing. The chloroplast genome of *Eucalyptus pauciflora* is 159,942 bp, contains 131 genes of known function, and confirms the phylogenetic position of *Eucalyptus pauciflora* as a close relative of *Eucalyptus regnans*.

**Conclusions:** Our results suggest that very accurate assemblies of chloroplast genomes can be achieved using a combination of at least 20x coverage of long- and short-reads respectively, provided that the long-reads contain at least ~5x coverage of reads longer than the inverted repeat region. We show that further increases in coverage give little or no improvement in accuracy, and that hybrid assemblies are more accurate than long-read-only or short-read-only assemblies.

## Background

Chloroplasts are important organelles in algal and plant cells, which generate carbohydrates by photosynthesis [1]. The chloroplast genome provides important information for phylogenetics, population-genetics and species identification [1–8], and is also the focus of genetic engineering because it contains many genes involved in photosynthesis [1].

The chloroplast genome is a double-stranded DNA molecule of around 120 kb - 160 kb in size in most plants, encoding around 100 protein coding genes [9, 10]. The chloroplast genome is usually circular, although some research suggests that it could be linear in certain developmental stages [11, 12]. The structure of chloroplast genome is highly conserved among plants, and usually consists of a long single copy and a short single copy region, separated by two identical inverted repeat regions. In some species, one copy of the inverted repeats has been lost during evolution [13]. The length of the inverted repeats usually ranges from 10 to 30 kb [9], although in extreme cases can be as short as 114 bp [14] or as long as 76 kb [15]. There are now more than 1500 chloroplast genomes available in the NCBI organelle genome database.

Accurate genome assembly is a key first step in the study of chloroplast genomes. Initial assemblies of the chloroplast genome relied on Sanger sequencing, which produces highly accurate reads of around 1 kb in length [16–18]. However, Sanger sequencing is expensive and time-consuming. Over the last decade, Sanger sequencing has been largely replaced by short-read sequencing as the primary approach to producing chloroplast genome assemblies [19–21]. Short-read sequencing produces large amounts of data for a relatively low cost, and has a high per-base accuracy. However, it produces DNA fragments that are typically around 50–400 bp in length [22], which can make genome assembly challenging.

A lack of long-range information can limit the accuracy of genome assemblies derived from short-read sequencing data. For example, the two inverted repeats can make it difficult to assemble the chloroplast genome into a single contig, because short-read data rarely contain sufficient long-range information to span an entire inverted repeat (~10–30 kb). Most studies which assembled chloroplast genomes from short-read data rely on assemblers designed for whole genome assembly, such as AbySS [23] and SOAPdenovo [24]. These assemblers produce multiple contigs, which were then assembled manually into a single contig according to the structure of existing chloroplast genomes, or by performing additional Sanger sequencing to confirm the conjunctions between the two single copy regions and the inverted repeats [19–21]. Assembling chloroplast genomes by performing synteny alignment with published chloroplast genomes may lead to inaccurate results if the chloroplast genome structure is not conserved (for example, the Chickpea chloroplast genome contains only one inverted repeat region [13]), or if the published chloroplast genome structure contains errors. And when Sanger sequencing is used to confirm an assembly, this removes some of the benefits of using short-reads to assemble chloroplast genomes.

New long-read sequencing technologies have the potential to allow for reference-free assembly of chloroplast genomes by combining some of the best features of Sanger and short-read sequencing. Like short-read technologies, long-read technologies produce large volumes of data for low cost. Technologies such as the MinION sequencing technology from Oxford Nanopore Technologies (ONT) and single-molecule real time sequencing technology from Pacific Biosciences (PacBio) routinely produce single reads longer than 10 kb [22, 25], and even up to 200 kb [25]. It is possible, therefore, for a single read to cover the entire chloroplast genome, or at least a very large section of it, suggesting that it should be feasible to use long-reads to perform reference-free assembly of the chloroplast genome. The main drawback of long-reads is that they have a relatively high per-base error rate of around 10–15% [22, 25], and ONT reads tend to systematically underestimate the length of homopolymer runs [26]. Typically, such errors may be mitigated by polishing the genome after assembly. Genome polishing involves remapping the reads back to the assembly to identify and fix single nucleotide and small structural errors [27, 28]. Chloroplast genome assembly using PacBio data has been reported [29–33], but no study has focused on employing ONT data to assemble chloroplast genomes. It is important to ascertain the best approach to assembling chloroplast genomes with ONT data, because it is increasingly widely used due to its high yield, low cost, and very rapid turnaround times from tissue to data [22, 34].

Hybrid assembly using a combination of long- and short-reads may be the best approach to assembling chloroplast genomes, because it can potentially combine the benefits of the length of long-reads and the accuracy of short-reads. However, we lack a comparison of the performance of long-read-only, short-read-only, and hybrid assembly approaches for the chloroplast genome. Furthermore, neither the minimum required length of long-reads, nor the optimum combination and coverage of long- and short-reads to obtain accurate chloroplast genome assemblies is known.

In this study, we set out to determine the best approach to chloroplast genome assembly using the chloroplast genome of *Eucalyptus pauciflora* (*E. pauciftora*) as a test case. Eucalypts are widely distributed in Australia, accounting for roughly 75% of the forest areas [35]. *E. pauciflora*, the snow gum, is an evergreen tree found in eastern Australia from close to sea level up to the tree line of the Australian Alps, displaying the broadest altitudinal range in the Eucalypts [36–38]. *E. pauciflora* is of particular interest due to its wide distribution and drought and cold tolerance [39–44]. However, no chloroplast genome for *E. pauciflora* is available. To evaluate the most efficient and accurate way to *de novo* assemble the chloroplast genome of *E. pauciflora*, we compared long-read-only assemblies, short-read-only assemblies and hybrid assemblies, with large ranges of different coverage and long-read lengths. We used Unicycler [45] to do the short-read-only and hybrid assemblies, because it is designed specifically for the assembly of small circular genomes. For long-read-only assemblies, we compared two assemblers: Hinge [46], an assembler designed for solving the long repeats problem in long-read assemblies of circular genomes; and Canu [47], one of the most popular long-read assemblers currently available. We also assessed the performance of a variety of read-correction methods, assembly software and polishing algorithms, and evaluated how the length of long-reads affects chloroplast genome assembly.

## Methods

The raw data have been submitted to NCBI under accession numbers SAMN09197243 (long-read data) and SAMN09197235 (short-read data). The final genome assembly of the E. pauciflora chloroplast genome, and the scripts necessary to reproduce the analyses in this study are available on github at https://github.com/asdcid/Chroloplast-genome-assembly. Subfolders in that repository are referenced throughout the methods where appropriate.

### Sample collection and DNA sequencing

#### Sample collection

Leaves were collected in March 2016 (for Illumina sequencing) and June 2017 (for MinlON sequencing) from the same branch of a single *E. pauciflora* individual near Thredbo, Kosciuszko Nation Park, New South Wales, Australia (Latitude −36.49433, Longitude 148.282983). Leaves were stored at 4°C until they were returned to the laboratory.

#### DNA extraction and quality control for Illumina sequencing

Leaves were freeze-dried, and total DNA was extracted using a CTAB protocol [48], then purified with a Zymo kit (Zymo Research Corp). TruSeq Nano libraries were constructed according to the manufacturer’s instructions and whole genome shotgun sequencing was carried out at the ACRF Biomolecular Resource Facility (the Australian National University, Canberra, Australia) on the Illumina HiSeq-2500 platform using 150 bp paired-end sequencing with a roughly 400 bp insert size. We used BBDuk v37.31 [49] for quality and adapter trimming of Illumina data, removing bases with a quality score <30 on the left or right side of a read. After trimming, reads shorter than 50 bp were removed, and only paired reads were kept. We used FastQC [50] to perform quality checks of libraries before and after trimming. Scripts for this analysis are available in the 1_pre_assembly/1_quality_control/short_read folder of the github repository.

#### DNA extraction and quality control for MinION sequencing

High molecular weight DNA was extracted from leaves using a protocol [51] based on Mayjonade’s method [52]. Libraries were prepared according to the ONT 1D ligation library protocol (SQK-LSK108). Long-read sequencing was carried out on a MinION sequencer using MinKNOW v1.7.3, and then basecalled using Albacore v2.0.2. We removed adapters from long-reads using Porechop v0.2.1 [53], and trimmed bases with quality <9 on both sides of reads using Nanofilt v1.2.0 [54]. We discarded reads shorter than 5 kb. Scripts for this analysis are available in the 1_pre_assembly/1_quality_control/long_read folder of the github repository.

### Chloroplast read extraction

In both the long- and short-read data, the majority of reads come from non-chloroplast sources, such as the nuclear genome, the mitochondrial genome and other contaminants, because we performed whole genome sequencing on DNA extracted from whole leaf tissues. To facilitate chloroplast genome assembly, we first extracted the chloroplast reads by attempting to align all reads to a dataset of 31 known *Eucalyptus* chloroplast genomes, which we refer to as the reference set (Table S1). The chloroplast genome is circular, but alignment algorithms rely on linear genomes. Simple linearization of the reference set would risk failing to capture reads that span the point at which the genomes were circularized. To avoid this, we duplicated and concatenated the sequence of each genome in the reference set. In this way, single reads that span the point at which the genome was circularized, or long-reads that span the entire chloroplast genome, would successfully map to the reference set. Short-reads were aligned to the reference set using Bowtie2 v2.2.6 [55] and long-reads were aligned to the reference set using Blasr v5.1 [56]. Scripts for this analysis are available in the 1_pre_assembly/2_cpDNAExtraction folder of the github repository.

### Genome assembly

#### Validation dataset preparation

As this is the first time that the *E. pauciflora* chloroplast genome has been sequenced, and because we want to assess and compare a range of different approaches to genome assembly, we used a subset of our short-read data to validate our genome assemblies. To do this, we randomly selected 100x coverage of paired-end Illumina reads (59,656 pairs of reads in total) from all of the chloroplast paired-end Illumina reads identified above. We excluded these reads from all genome assemblies, and instead used them exclusively to assess and compare the genome assemblies we produced. In general, we should expect that reads in the validation set will map more successfully to chloroplast genomes that are closer to the true genome from which the reads were sequenced. Because of this, we can use the mapping rate and the error rate of mapped validation reads to different genome assemblies to compare different genome assemblies. The mapping rate may not reach 100% even with a perfect genome assembly, because we cannot guarantee that all of reads in the validation set come from the chloroplast genome (e.g. some may come from nuclear copies of chloroplast genes, or from contaminants) and some reads will also contain sequencing errors. Nevertheless, as long as the vast majority of the validation reads come from the chloroplast genome of *E. pauciflora*, it should be the case that the mapping rate of the validation reads will increase monotonically as the accuracy of the chloroplast genome assembly increases. We also expect the error rate of mapped validation reads to decrease monotonically as the accuracy of the genome assembly increases. Short Illumina reads typically have a low error rate of ~0.0010 errors per base [57], therefore for a perfect genome assembly we might expect the mapped validation reads to have an error rate of at most ~0.0010 errors per base (the true number is likely to be lower than this, because we filtered out reads and bases with low quality scores before constructing the validation set). Assembly errors will increase the error rate in the mapped validation reads. Thus, if the error rate of the mapped reads is greater than ~0.0010 errors per base, this suggests that there are errors that cannot be attributed to sequencing error, and are likely to represent assembly errors. If the error rate is less than 0.0010 errors per base, this suggests there are few if any errors in the assembly, although it does not guarantee that the assembly is error free. Scripts for random read selection are available in the 2_assembly/randomSelection folder of the github repository.

#### Long-read-only assembly

We performed 24 long-read-only assemblies, using all combinations of two assemblers, Hinge and Canu, and 12 different coverages (calculated assuming a 160 kb genome size): 5x, 8x, 10x, 20x, 40x, 60x, 80x, 100x, 200x, 300x, 400x, 500x. Reads for each coverage were selected randomly from the full set of filtered and trimmed long-reads. The assemblers were run with the default settings, except: (i) in Hinge, the read filter cut-off was set to 10, as it failed to assemble reads with default setting; and (ii) in Canu, the correctErrorRate was increased to 0.154 when the coverage was less than 40x according to the recommendation in the Canu manual. Hinge failed to assemble the chloroplast genome with 5x, 8x and 10x long-read coverage. Scripts for this analysis are available in the 2_assembly/longReadOnly folder of the github repository.

#### Short-read-only assembly

We performed 48 short-read-only assemblies using default settings in Unicycler v0.3.1. These 48 assemblies comprised 12 different coverages (5x, 8x, 10x, 20x, 40x, 60x, 80x, 100x, 200x, 300x, 400x, 500x), each combined with four types of short-read error correction: (i) no correction; (ii) SPAdes correction [58]; (iii) Karect correction [59]; and (iv) Karect+SPAdes correction. A recent study [60] suggested that Karect performs better than other error correction methods, while SPAdes is the correction method built into the Unicycler pipeline. Corrected and uncorrected paired-end reads for the different coverages were randomly selected from the total set of filtered and trimmed short-reads prior to assembly. Scripts for this analysis are available in the 2_assembly/shortReadOnly folder of the github repository.

#### Hybrid assembly

We performed 576 hybrid assemblies combining long- and short-reads. All assemblies were performed using default settings in Unicycler v0.3.1, except for those assemblies which used only Karect to correct the reads, for which we turned off the SPAdes correction in Unicycler). The 576 assemblies comprise all 144 possible combinations of 12 different coverages of both long- and short-read used above (5x, 8x, 10x, 20x, 40x, 60x, 80x, 100x, 200x, 300x, 400x, 500x), repeated for four types of short-read error correction: (i) no correction; (ii) SPAdes correction; (iii) Karect correction; and (iv) Karect+SPAdes correction. We did not perform long-read error correction prior to assembly, because Unicycler is designed to work with raw long-read data, and we polish all genome assemblies using the best available short-read polishing tools. Scripts for this analysis are available in the 2_assembly/hybrid folder of the github repository.

### Hybrid assembly with different lengths of long-reads

To investigate how the length of long-reads affects the accuracy of chloroplast genome assemblies, we performed 576 genome assemblies that combined short-reads with long-reads of different lengths. All assemblies were performed using Unicycler v0.3.1. The 576 assemblies comprise all 96 possible combinations of short- (5x, 8x, 10x, 20x, 40x, 60x, 80x, 100x, 200x, 300x, 400x, 500x) and long-read coverage (5x, 8x, 10x, 20x, 40x, 60x, 80x, 100x), repeated for six different length categories for the long-reads: 5–10 kb, 10–20 kb, 20–30 kb, 30–40 kb, 40–50 kb, 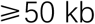. As above, sets of reads were picked randomly from each length specific sub-dataset. Short-reads were corrected by Karect, and long-reads were uncorrected, as above. We did not investigate higher coverage of long-reads because 100x was the maximum coverage category available for the longest subset of long-reads 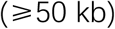. Assembly scripts for this analysis are the same as for the hybrid assemblies, except for the input files.

#### Genome polishing

To facilitate certain comparisons among genome assemblies, we manually assembled the multiple contigs into a single contig from some genome assemblies. For example, the long-read-only assemblies, short-read-only assemblies, and hybrid assemblies with <20x long-read coverage all tended to produce multiple contigs, occasionally with some regions of the chloroplast genome represented more than once. This is, of course, useful information for comparing assembly performance. However, in order to calculate comparable estimates of the per-base error rate from our validation set, we manually removed duplicated regions from these assemblies and then manually combined all remaining contigs together to create a single contig spanning as much of the chloroplast genome as possible. To do this, for assemblies with multiple contigs, we used *E. regnans* (NCBI accession NC_022386.1) as a reference, which is a close phylogenetic relative of *E. pauciflora*, and created pairwise genome alignments between this reference and the contigs from each assembly using Mummer v3.23 [61], after removing contigs shorter than 5 kb. We then used a python script to remove duplicate regions according to the pairwise genome alignments. Scripts for this analysis are available in the 3_post_assembly/1_same_structure folder of the github repository.

All genome assemblies were polished at the individual base level using the best available tools. Hybrid and short-read-only assemblies were polished using Pilon v1.22 [62], and long-read-only assemblies were polished using Racon [28], Nanopolish v0.8.1 [63], and Racon + Nanopolish v0.8.1. Pilon uses information in short-reads to polish out errors from assemblies, whereas Racon and Nanopolish use information in long-reads. Nanopolish is designed for ONT data, but Racon is designed for ONT and PacBio data. We ran Nanopolish and Pilon for multiple iterations until the polished genome remained unchanged. We ran Racon for ten iterations, because the polished genome continued changing by tiny amounts after 10 iterations during testing. Scripts for this analysis are available in the 3_post_assembly/2_polish folder of the github repository.

### Evaluation of different assembly performance

A perfect genome assembly would cover the entire genome in a single contig, with the maximum possible percentage of validation reads mapping successfully with the lowest possible error rate, given the limitations of the accuracy of the sequencing platform and the mapping software. Therefore, to assess and compare our genome assemblies, we considered five key statistics: (i) the number of contigs output by the raw assembly; (ii) the average per-base coverage of the reference genome by the contigs (calculated by the sum of any part of any contig that aligns successfully to the *E. regnans* chloroplast genome divided by the length of the *E. regnans* chloroplast genome), which should equal ~1.0 for a perfect assembly but may be much higher than one for assemblies that contain duplicated regions; (iii) the sum of the length of all contigs output by the assembler; (iv) the percentage of the validation reads successfully mapped to the polished genome; (v) the error rate of the validation reads that successfully mapped to the polished genome. As mentioned above, the error rate here is the sum of mismatches and indels (insertions or deletions) between the validation reads and the assemblies, because it is not possible to know in advance whether a mismatch or indel between a read and an assembly derives from an error in the read or the assembly (although this may be determined post-hoc by analysis of the aligned reads to the assembly, see below). We mapped the validation reads to each genome with default settings in Bowtie v2.2.6, and extracted the mapping rate and general error rate from the result using Qualimap v2.2.1 [64]. Scripts for this analysis are available in the 3_post_assembly/3_assembly_quality_control folder of the github repository.

### Final genome assessment

To further assess the assembly with lowest error rate, highest mapping rate, and coverage closest to 1.0, we attempted to estimate the number of possible errors and heteroplasmic sites in that assembly using all of the long- and short-read data available to us. To do this, we mapped all of the short-reads with Bowtie v2.2.6, and long-reads using Ngmlr [65] (because Blasr failed to produce BAM file for ONT data) to the assembly after duplicating and concatenating it as above, and compared the mapped reads at each site by visualising the alignments in IGV [66], and by using samtools v1.5 [67] and varScan v2.4.0 [68] to look for mismatches and indels between the short-reads and the assembly, and Nanopolish v0.8.1 to look for mismatches and indels between the long-reads and the assembly. We considered that a true assembly error would be likely to produce consistent mismatches or indels between the assembly and both the long- and short-reads. Both technologies have their limitations which may produce well-supported mismatches or indels between the assembly and one read type. For example, short-reads that derive from regions of the nuclear or mitochondrial genome that are similar to the chloroplast genome may map with high probability to the chloroplast genome, but this mapping is far less likely to occur with long-reads, which in our dataset are a minimum of 5 kb. Similarly, ONT reads are known to contain biases with respect to the length of homopolymer regions, but this bias should not occur in short-reads. Thus, if the frequency of the non-reference base of a mismatch or indel is similar between the short- and long-read mappings, then this is most likely to indicate either an assembly error or a heteroplasmic site. Scripts for this analysis are available in the 3_post_assembly/4_SNP_call folder of the github repository

### Gene annotation

We used GeSeq [69] to annotate the best assembly, followed by ARAGORN v1.2.38 [70] to annotate the tRNAs. Since GeSeq failed to detect the very short exons, genes containing such exons, such as *petB* and *petD*, were manually annotated based on the annotations in the reference set of chloroplast genomes.

### Phylogenetic analysis

We added the *E. pauciflora* chloroplast genome estimated here to the reference set of chloroplast genomes used above, giving a dataset of 32 complete and well-annotated chloroplast genomes. We split each genome into 312 fragments based on their annotations, where each fragment represents a single exonic or non-exonic region. We then produced 312 individual alignments of the 32 sequences (one alignment for each fragment) using Clustal Omega v1.2.4 [71]. We then manually examined the 312 alignments, deleted regions of each alignment that could not be confidently aligned, and manually adjusted the remaining regions. Subsequently, we concatenated the 312 alignments to create a single alignment with 32 sequences and 156,245 bases. We estimated a phylogeny from this alignment using IQ-TREE v1.5.5 [72], and ModelFinder [73] with default settings, and 1000 ultrafast bootstrap replicates [74]. Scripts for this analysis are available in the phylogenetic_analysis folder of the github repository.

## Results

### Sequencing

We generated 6.00 Gb of raw long-read data, comprising 705,554 reads with a mean length of 8,504 bp. We generated 3.19 Gb of short-read data, comprising 21,114,786 150 bp paired-end reads. After trimming adapters and low-quality bases, and mapping the reads to the reference set (Table S1), we recovered 0.57 Gb of long-read data comprising 28,777 reads with a mean length of 19,807 bp (minimum 5,002 bp and maximum 150,181 bp), and 0.38 Gb of short-read data comprising 2,840,345 paired-end reads. Assuming that the *E. pauciflora* chloroplast genome size is ~160 kb, this represents a total coverage of ~3,600x for long-reads and ~2,400x for short-reads.

### Genome assembly comparison

Of the 648 genome assemblies we performed, the best genome assemblies were the hybrid assemblies with at least 20x coverage of both long- and short-reads (Figure 1). Hybrid assemblies with lower coverage of either data type, and assemblies that relied on only one data type, were of lower quality. In the following, we discussed each of the assembly categories in more detail.

**Fig 1.**
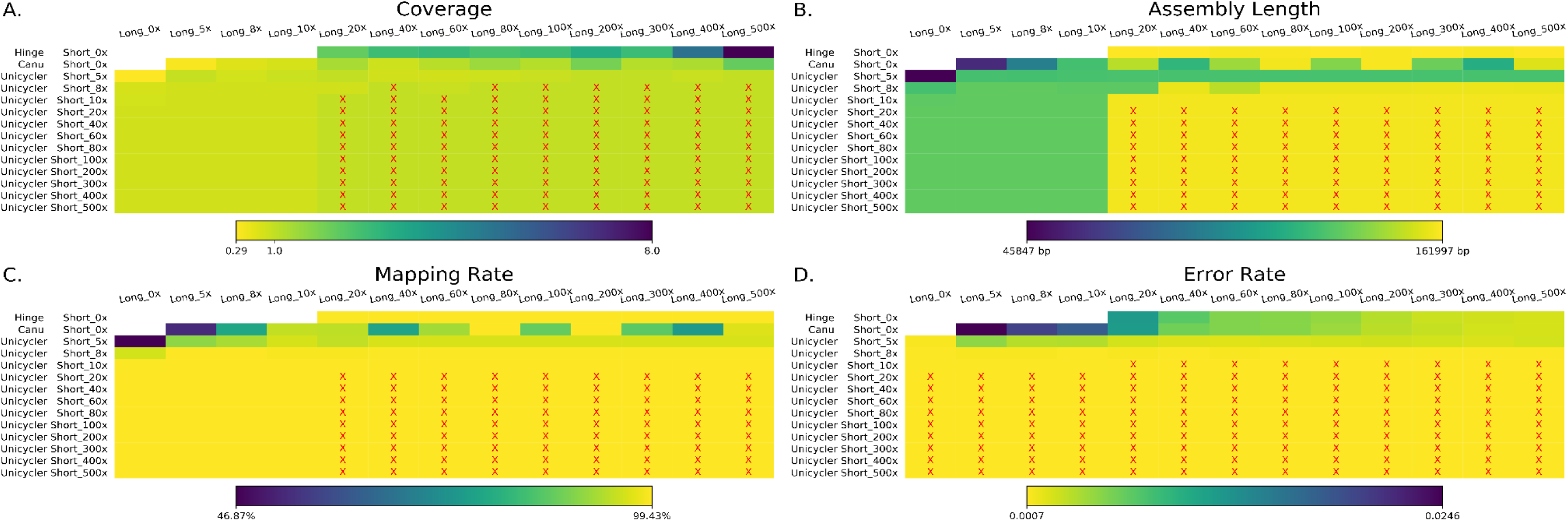
Comparison of chloroplast genome assemblies. The coverage of the long- and short-reads is shown along the top and left-hand-side of each panel, respectively. The left-hand-side also shows which assembler was used for each row of assemblies in that panel. Hinge and Canu are long-read-only assemblers, whereas Unicycler is short-read-only and hybrid assembler. The Hinge and Canu results in B, C, D were polished by Racon+Nanopolish, and the Unicycler results used Karect-corrected short-reads. **A.** The total coverage of the chloroplast genome across all contigs output by the assembler. Panels marked with a red ‘x’ contained a single contig covering the whole chloroplast genome. The heatmap indicates the chloroplast genome coverage. **B.** The assembly length of different assemblies after manual curation (e.g. removing duplicate regions). Panels marked with an ‘x’ denote assemblies wit h the expected length, in the range 155,938 bp-155,945 bp. **C.** The mapping rate of validation reads to the assemblies after manual curation. Assemblies with highest mapping rate (99.43%) are marked with a red ‘x’. **D.** The average per-base error rate of validation reads mapped to each manually-curated genome assembly. Assemblies with the lowest error rate (0.0007) are marked with a red ‘x’.

#### Long-read-only assembly

For long-read-only assemblies, the best assembly was produced by Hinge with 500x long-read coverage and polishing using Racon + Nanopolish (Figure S1). Neither Hinge nor Canu were able to assemble the entire chloroplast genome into a single contig (Figure S1a), so all long-read-only assemblies had to be manually curated (see Methods). In general, Canu assembled one or two contigs which failed to cover some regions of the genome or covered some regions twice, whereas Hinge assembled at least two contigs containing many duplicated regions. Canu produced assemblies from all 12 different long-read coverages (5–500x; the second row of Figure 1a), but only the assemblies generated from 80x and 200x coverage had a total length that was close to the expected 160 kb after manual curation (Figures 1b and S1b). Surprisingly, the total length of the Canu assemblies varied widely with relatively minor changes in coverage (E.g. ~145 kb at 60x, ~160 kb at 80x, and ~140 kb at 100x coverage; Figures 1b and S1b). Hinge failed to produce any contigs when the coverage was lower than 20x (top row of Figure 1a), but produced contigs that could be manually curated into near complete chloroplast genomes (with lengths very close to 160 kb) when the coverage was 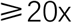 (Figures 1b and S1b).

Validation of the long-read-only assemblies after manual curation showed that the assemblies produced by Hinge were more accurate than those produced by Canu (Figure 1c). This is because Hinge was able to assemble contigs covering the entire chloroplast genome. The error rate of mapped validation reads decreased predictably as the long-read coverage increased for both Canu and Hinge assemblies (Figures 1d and S1d). For the Hinge assemblies, the mapping rate of the validation reads increased predictably with the increased coverage of input reads, up to a maximum of 99.41% with 300x or 500x input read coverage (Figures 1c and S1c). For the Canu assemblies, the mapping rate of the validation reads was the highest (~99.40%) at 80x and 200x input read coverage (Figures 1c and S1c), but varied substantially at lower coverage, reflecting the fact that many of the Canu assemblies were missing large portions of the chloroplast genome. These data suggest that there is a complex relationship between assembly accuracy and input read coverage when assembling chloroplast genomes with ONT data using Canu.

Polishing the long-read-only genome assemblies with both Racon and Nanopolish resulted in more accurate assemblies than polishing them with either Racon or Nanopolish alone (Figures S1c and S1d). For Canu assemblies, when polishing with Racon alone, assembly accuracy improved from an error rate (mismatches and indels) of ~0.0300 per base of the validation reads to ~0.0060 per base as coverage increased up to ~100x, but did not improve further at higher coverage. When polishing with Nanopolish or Racon+Nanopolish, assembly accuracy continued to improve up the maximum coverage we examined (500x; Figures S1c and S1d), at which the error rate was 0.0022 per base of the validation reads in the best assemblies. The situation was the same in Hinge assemblies. Since the expected error rate of the validation reads likely to be at most 0.0010 (see above), this suggests that at least 0.0012 mismatches or indels per base in the validation reads mapped to the long-read-only assemblies comes from errors in the assemblies themselves.

#### Short-read correction

For short-read-only and hybrid assemblies, we compared the effect of four different approaches to short-read correction: no correction, Karect, SPAdes, and Karect+SPAdes. When the short-read coverage was 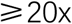, we found no meaningful differences between these four conditions (Figures S2-S5). When the short-read coverage was lower than 20x, we found that correction with Karect gave slightly higher genome assembly accuracy than the other three conditions (i.e. slightly higher mapping rates and lower per-base error rates with the validation reads; Figures S2-S5). Therefore, we focus on comparing short-read-only and hybrid assemblies in which short-reads were corrected using Karect.

#### Short-read-only assemblies

Short-read-only assemblies with coverage 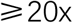 produced three complete contigs corresponding to the three major structural regions of the chloroplast genome: the long single copy, the short single copy and the inverted repeat (Figure S2a). After manual curation (see Methods), in which we assembled the resulting three contigs by hand and polishing, the short-read-only assemblies had high mapping rates of the validation reads (99.36%) and low rates of mismatches and indels between the validation reads and the assemblies (0.0007). This error rate is lower than the expected error rate of the validation reads, suggesting that the short-read-only assemblies may contain few or no errors. One small source of error in the short-read-only assemblies, which is not reflected in these statistics, is that in these assemblies ~10 bp from the beginning or end of long single copy or short single copy region are sometimes incorrectly assigned to the inverted repeat region, vice versa. This may cause minor errors in assemblies when whole chloroplast genomes are manually assembled from the three contigs that result from short-read-only assemblies.

#### Hybrid assemblies

Hybrid assembly with Unicycler performed better than all other assemblies as long as both long- and short-read coverage was at least 20x (Figures 1 and S2). In these cases, the entire chloroplast genome was assembled into a single contig without manual curation, and the validation reads showed a mapping rate of 99.43% and an error rate of 0.0007 after polishing. As above, this suggests that these hybrid assemblies likely contained few or no errors. When the long-read coverage was less than 20x, the long-reads made no meaningful difference to the hybrid assemblies when compared to short-read-only assemblies. When coverage of long-reads was 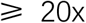, all assemblies with the same short-read coverage were exactly the same. These results suggest that the long-read information is useful for confirming the correct conjunctions between the two single copy regions and the inverted repeats in Unicycler. Assemblies using only 5x short-read coverage were missing ~40 kb of the chloroplast genome, regardless of the long-read coverage (Figure S2b). The length of hybrid assemblies with 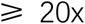 coverage of both long- and short-reads were all very similar, differing by at most four bases from a length of 159,942 bp.

### Hybrid assembly with different lengths of long-reads

Hybrid assembly performance was highly dependent on the length of the long-reads. Hybrid assembly produced a single contig spanning the entire complete chloroplast genome with as little as 5x long-read coverage and 8x short-read coverage, provided that all the long-reads were longer than 30 kb (Table 1 and Figures S6-S11), which is long enough to cover the entire inverted repeat region (~26 kb). When long-reads were shorter than 20 kb, the hybrid assembly failed to assemble the entire chloroplast genome into a single contig (Table 1, Figures S6 and S7), likely because these reads are shorter than the inverted repeat region. When long-reads were 20–30 kb, hybrid assembly produced a complete assembly with a single contig with at least 20x long-read coverage and at least 10x short-read coverage.

**Table 1.** The minimum coverage of long-/short-reads required to assemble one contig spanning the entire chloroplast genome.

| Long-read length range | 5-10 kb | 10-20 kb | 20-30 kb | 30-40 kb | 40-50 kb | ≥ 50 kb |
| --- | --- | --- | --- | --- | --- | --- |
| Long-read | N/A | N/A | 20x | 5x | 5x | 5x |
| Short-read | N/A | N/A | 10x | 8x | 8x | 8x |

N/A: No one contig spanning the entire chloroplast genome can be assembled under this long-read length range.

### Final genome assessment

To choose one assembly as the *E. pauciflora* chloroplast reference genome, we compared all hybrid assemblies with 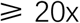 long- and short-read coverage. The 81 assemblies fall into six groups of identical assemblies (Figure 2), all of which have identical mapping and error rates (Figure 1c and 1d). All assemblies with the same short-read coverage were identical, regardless of their long-read coverage. Assemblies with 40x, 200x or 500x short-read coverage were identical, as were assemblies with 300x and 400x short-read coverage, while assemblies with 20x, 60x, 80x and 100x short-read coverage were all unique. The differences between these six groups of assemblies fall into just three regions of the chloroplast genome (Figure 2): (i) an adenine homopolymer site, 9048–9061, containing six variants; (ii) an adenine homopolymer site, 32043–32059, in which the 80x short-read coverage assemblies have a small deletion; (iii) a thymine homopolymer site, 51964–51975, in which the 20x short-read coverage assemblies have a short deletion.

**Fig 2.**
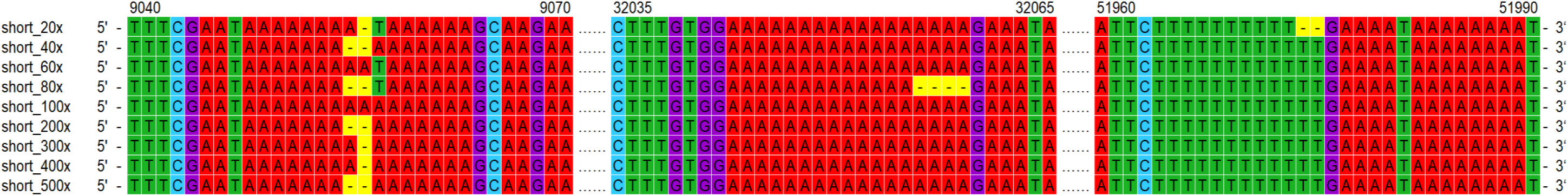
Comparison of hybrid assembly sequences with 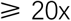 long- and short-read coverage. Short_20x indicates 20x coverage of short-reads were used in these assemblies (with 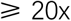 long-read coverage). For hybrid assemblies with 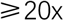 short-reads, if the long-read coverage was 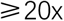, all assemblies with the same short-read coverage was identical. The number at the top is the position.

To distinguish which variant in each of these regions was most likely to be correct, we mapped all of the long- and short-reads to all six assemblies. This revealed that the identical assemblies from the 40x, 200x, and 500x short-read coverage datasets are most likely to be correct. In the first adenine homopolymer site, both short- and long-reads rejected the thymine substitution (which is present in the 20x, 60x, and 80x short-read coverage assemblies, Figure 2), but neither the short- nor the long-reads provided clear preference for the length of the homopolymer. The long-reads provided little useful information due to their high rate of systematic error around homopolymers. Roughly the same number of reads map successfully to this region regardless of whether the assembly has 14, 15, or 16 adenines in this homopolymer, and in all cases no roughly 10% of short-reads disagree with the length of the homopolymer in the assembly. This may be the result of sequencing error, or mapping error, or heteroplasmies affecting this site. For the other two homopolymer sites, the short-reads clearly showed that the deletions in the 20x and 80x coverage assemblies in these regions where likely to be assembly errors (>50% of short-reads show insertions in both regions). Based on these observations, we selected the genome assembly derived from 500x coverage of both short- and long-read data as the final genome assembly. Despite the uncertainty in the homopolymer at position 9048–9061, we preferred this assembly because it is derived from the most data.

The mapping results suggest that the per-base accuracy of the final genome assembly is very high, and is unlikely to contain any errors. We identified all sites that show a non-reference base or small indel at a frequency of 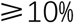, revealing 1016 sites for the mapped long-reads and one for the mapped short-reads. Of these, there are no sites that contained a non-reference allele at 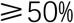 frequency in either the long- or the short-read mapping. This suggests that the final genome assembly is unlikely to contain any errors. The high number of sites with a non-reference allele at 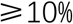 for the mapped long-reads is expected given their much higher error rate, and biases associated with the estimation of the length of homopolymer runs. The one site identified with a non-reference base 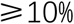 in the short-reads was site 51961 in which 47.71% of bases support a T insertion. This insertion is located in a homopolymer T region (Figure S12), and so the ONT long-reads can provide little information on whether the insertion is likely to be an assembly error, a true variant (e.g. due to heteroplasmy), sequencing error in the short-reads, or a mapping error (e.g. due to a proportion of short-reads being derived from DNA transferred from the chloroplast to the mitochondrial or nuclear genomes [75]).

### *E pauciflora* chloroplast genome annotation, and phylogenetic analysis

The *E. pauciflora* chloroplast genome is 159,942 bp in size, comprised of two inverted repeat regions of 26,367 bp, a long single copy region of 88,787 bp, and a short single copy region of 18,421 bp. We identified and annotated 131 genes of known function including 37 transfer RNA genes and 8 ribosomal RNA genes (Figure 3a). All ribosomal RNA genes are located in the inverted repeat regions. Five genes, *psbL, infA, ycf1* and two copies of *ycf15*, were annotated as pseudogenes.

**Fig 3.**
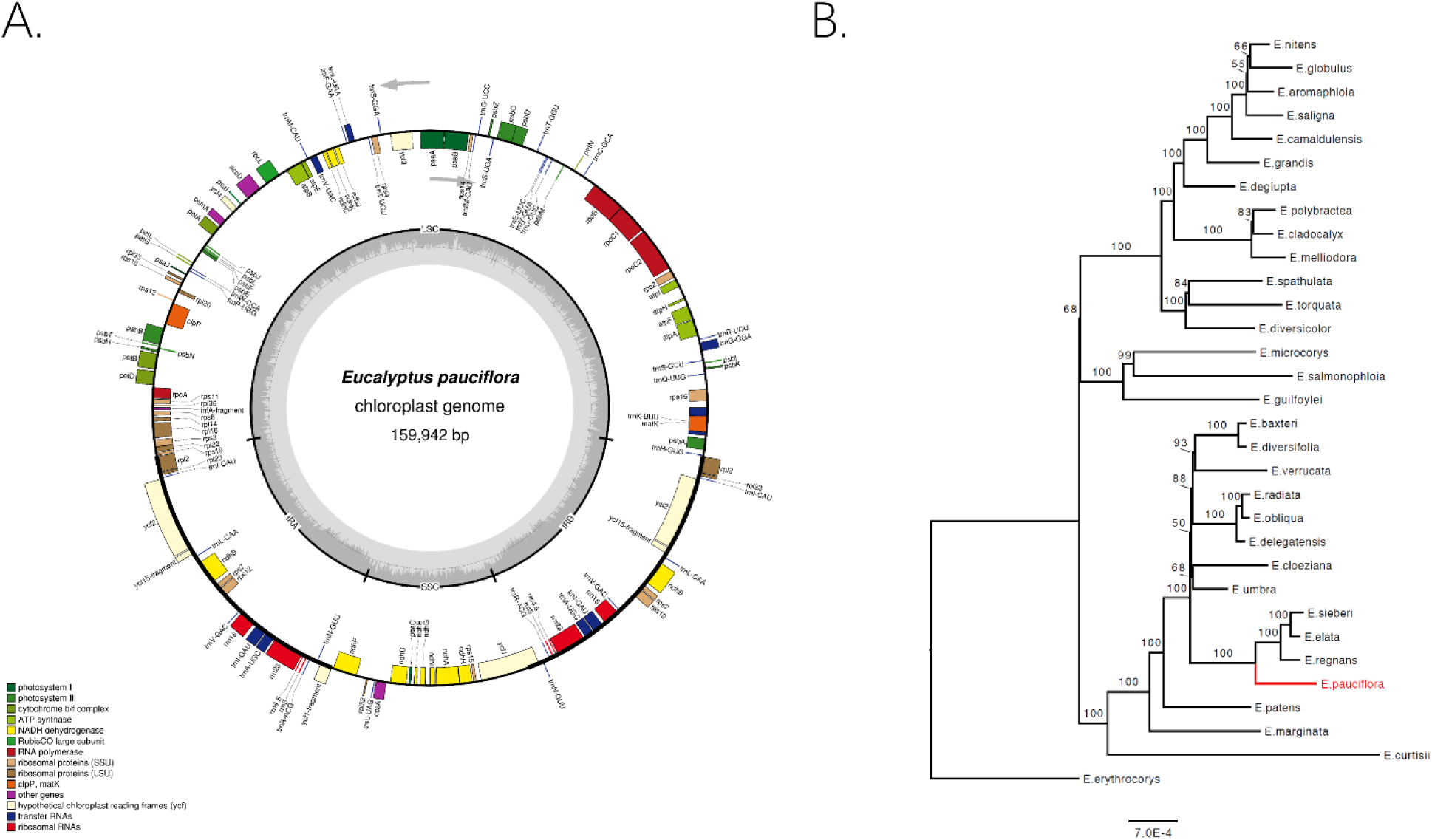
**A.** Annotated *E. pauciflora* chloroplast genome. Genes shown on the inside of the circle are transcribed clockwise, whereas genes shown on the outside of the circle transcribed counterclockwise. The grey region in the inside circle shows the GC content across the chloroplast genome. This figure was produced by OGDraw v1.2 [88]. **B.** A phylogenetic tree of 32 *Eucalyptus* taxa based on analysis of full chloroplast genomes.

The phylogenetic analysis places *E. pauciflora* as a sister to a clade comprised of *E. regnans, E. elata*, and *E. sieberi* with high bootstrap support (Figure 3b). As expected, interrelationships among the other 31 *Eucalyptus* chloroplast genomes are the same as reported previously [5]. The placement of *E. pauciflora* is consistent with previous *Eucalyptus* analyses using Diversity Array Technology markers [76].

## Discussion

By comparing a large range of different approaches to chloroplast genome assembly, we show that hybrid assembly with at least 20x coverage of long-reads (containing at least 5x coverage of reads large than the inverted repeat region) and 20x coverage of short-reads is sufficient to assemble the entire chloroplast genome into a single contig with few or no errors. We also show that very similar accuracy can be obtained from short-read-only assemblies, although in this case the genome is assembled into three contigs representing the three main regions of the chloroplast genome. Long-read-only assemblies using ONT data produced multiple contigs and contained many errors, even after polishing with long-read polishing algorithms. We demonstrate how multiple genome assemblies from the same data can be rigorously compared by retaining a subset of reads as a validation set. Finally, by re-mapping both our long- and short-reads to our final assembly, we show that it is likely that our final assembly has few or no errors, and identify one sites in which the lengths of homopolymers were difficult to determine, even with very high coverage of long-and short-reads.

Short-read-only assemblies of the chloroplast genome were highly accurate, but were divided into the three regions of the chloroplast genome - the long single copy, short single copy, and inverted repeat. Such assemblies have two limitations. First, they must be manually combined into a full-length chloroplast genome, and this relies on comparison to a reference genome. Second, if using Unicycler as the assembler, although it can assemble the three regions of chloroplast genome, it occasionally placed ~10 bp from the end of one region onto the start of another, such that the accuracy of manually-curated short-read-only assemblies may be rather low at the junctions between regions. This may not matter for some approaches (e.g. phylogenetics, in which such regions could be safely ignored), but may be important for other applications (e.g. population or functional genetics in which variants in these regions may be the focus of a study).

Our study suggests that there are two key advantages to hybrid assemblies compared to short-read-only assemblies. First, the addition of as little as 5x coverage of long-reads of sufficient length that span the inverted repeat (roughly 10–30 kb in most species) results in assembling the entire chloroplast genome into a single contig. This will be beneficial for those who wish to obtain a reference-free estimate of the overall structure of the chloroplast genome (e.g. for researchers interested in structural variation [77–80]), and for those who wish to accurately infer the sequence of the junctions between the major regions of the genome without additional Sanger sequencing. Second, long-reads provide useful information for assessing the assembly errors or heteroplasmies in chloroplast genome assembly. The limited length of short-reads (150 bp in this study) will increase the rate at which these reads map incorrectly, particularly given the existing copies of sections of the chloroplast genome in the mitochondrial or nuclear genome [81–84]. This problem should be much reduced for long-reads, which in our study had a minimum length of 5 kb.

Our results differ from those reported in a recent study of long-read-only and short-read-only chloroplast genome assemblies, which suggested that there were substantial benefits of long-read-only assembly for chloroplast genomes assembly [29]. That study used PacBio data (we used ONT data here), and compared the resulting assemblies to those generated from Illumina data (as we used here). That study suggested that long-reads were beneficial based on the observation that short-read-only assemblies recovered just ~90% of the chloroplast genome assembled into seven contigs, with a relatively high proportion (0.12%) of uncertain sites. We were able to successfully assemble 100% of the chloroplast genome into three contigs (corresponding directly to the three major structural regions of the chloroplast genome) with very high accuracy, given just 20x coverage of short-read data. The difference between these two studies is likely due to recent improvements in genome-assembly algorithms, and in particular the development of Unicycler, which is designed specifically for the assembly of circular genomes from short-read and combined short- and long-read data. These improved genome assembly algorithms mitigate many of the previous limitations of short-read-only assemblies of the chloroplast genome, making the differences between approaches less severe than they were previously.

Assembling highly accurate chloroplast genomes from purely long-reads generated on the ONT platform remains challenging. The long-read-only assemblers we assessed were unable to assemble the genome into a single contig, although they were able to assemble multiple contigs which could subsequently be manually curated to produce full-length chloroplast genomes (Figures 1 and S1). However, the single-base accuracy of these genome assemblies (after polishing) remained much higher than those of short-read-only or hybrid assemblies (after polishing). Long-read-only assemblies are likely to be more accurate at the single base level when using PacBio sequencing, because the error profile of PacBio sequencing is less systematically biased than that of ONT sequencing. However, in agreement with our findings, a recent study showed using PacBio sequencing data it was not yet possible to assemble the entire chloroplast genome into a single contig without post-assembly processing [29–31, 85].

## Conclusions

Our results show that it is possible to produce complete and highly-accurate chloroplast genome assemblies by combining at least 20x coverage of long- and short-read data, respectively. Given the low cost and simplicity of generating long-read data from the ONT sequencer (MinION) [22, 34], the extremely low cost of producing high-coverage short-read data using Illumina technology, and the potential to multiplex multiple samples on both devices, this provides a clear path towards producing multiple highly-accurate and complete chloroplast genome assemblies for very low cost. The ability to recover the genome in a single contig will also be of great benefit to those interested in assembling chloroplast genomes with atypical structures.

Perhaps the biggest remaining challenge with the hybrid assembly approach is the production of long-reads that span the entire inverted repeat region. We showed that it is necessary to include at least 5x coverage of such reads to gain the benefits of the hybrid assembly approach. In most species, the inverted repeat region is ~10–30 kb, and successfully extracting and sequencing DNA containing sufficient intact molecules longer than 30 kb remains difficult.

Nevertheless, many groups are now working on establishing high-molecular-weight DNA extraction protocols (e.g. [86, 87]), suggesting that the remaining issues are unlikely to be insurmountable for most species.

Broadly speaking, our analyses may also provide hints on obtaining high-quality assembly of other circular genomes, such as those of other organelles and bacteria.

## Abbreviations

bp: base pair
kb: kilo base pair
Gb: giga base pair
ONT: Oxford Nanopore technologies
PacBio: Pacific Biosciences
NCBI: National Center for Biotechnology Information

## Declarations

### Acknowledgements

We thank Eleanor Beavan for help with tissue collection and DNA extractions.

### Funding

This research is supported by the Australian Research Council Future Fellowship, FT140100843.

### Availability of data and materials

The datasets supporting the results of this article are available on NCBI, https://www.ncbi.nlm.nih.gov/ (accession number: MG921592), and github, https://github.com/asdcid/Chroloplast-genome-assembly.

### Authors’ contributions

AS and RL carried out sample collection and AS performed DNA extraction for Illumina sequencing. RL and MS carried out sample collection for MinlON sequencing. MS and BS performed DNA extraction, library preparation, and MinION sequencing. WW, DK, RL, MS, and BS were involved in data QC and discussion. WW and RL planned all analyses. WW conducted all data analyses and drafted the original manuscript. RL and WW finished the manuscript. All authors read and approved the final manuscript.

### Ethics approval and consent to participate

Not applicable.

### Consent for publication

Not applicable.

### Competing interests

The authors declare that they have no competing interests.

## Additional files

**Figure S1.**
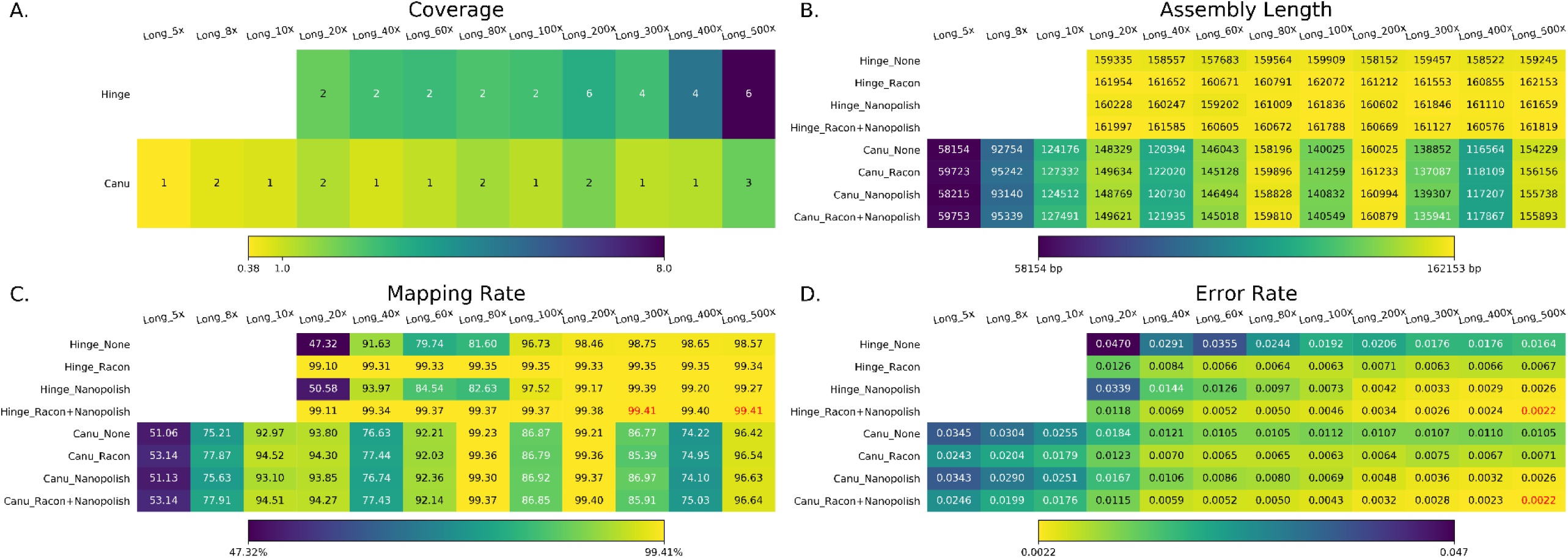
The summary of long-read-only assemblies. Long_5x indicates the 5x coverage of long-read was used in the assembly. Hinge failed to assemble genome with <20x coverage. None, Racon, Nanopolish and Racon+Nanopolish mean the different genome polishing pipeline. **A.** The total coverage of the chloroplast genome across all contigs output by the assembler. The number is the number of contigs of each assembly, whereas the heatmap is the genome coverage (it could be over 100% if some duplications exist). **B.** The assembly length of different assemblies after manual curation (e.g. removing duplicate regions). **C.** The mapping rate of validation reads to the assemblies after manual curation. **D.** The average per-base error rate of validation reads mapped to each manually-curated genome assembly.

**Figure S2.**
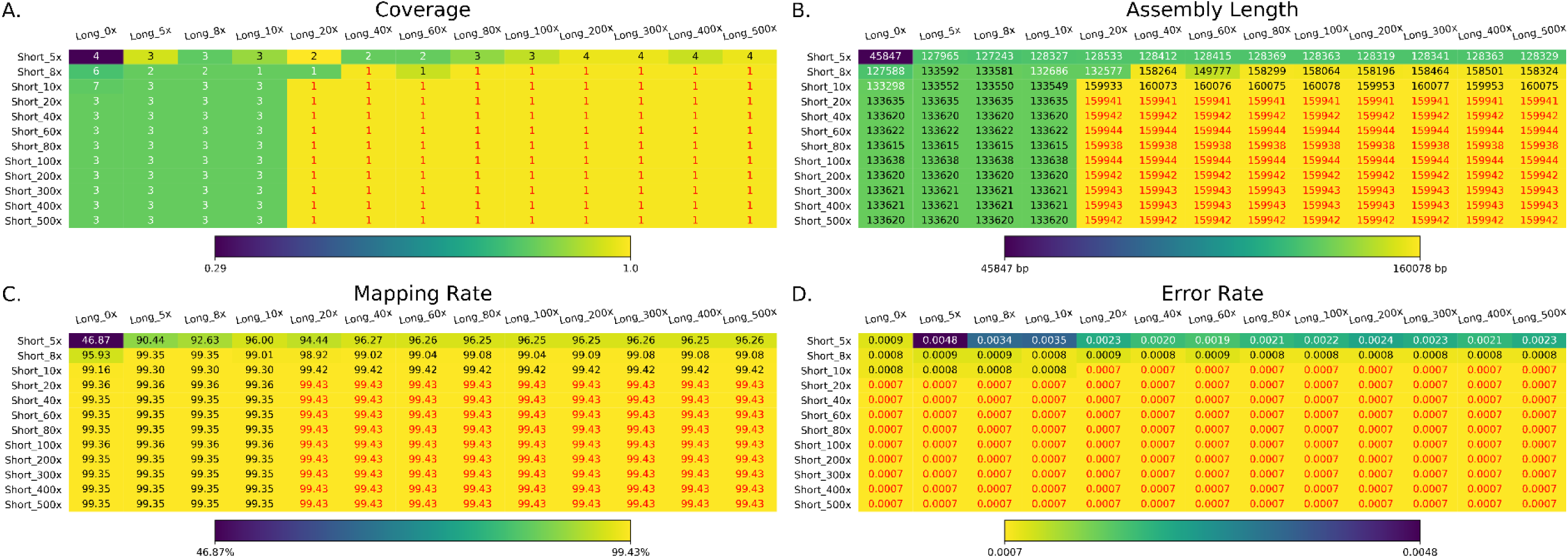
The summary of short-read-only and hybrid assemblies with Karect short-read correction. Long and Short indicate the coverage of this assembly. **A.** The total coverage of the chloroplast genome across all contigs output by the assembler. The number is the number of contigs of each assembly, whereas the heatmap is the genome coverage (it could be over 100% if some duplications exist). Numbers marked with red contained a single contig covering the whole chloroplast genome. The heatmap color is reversed compared to the Figure 1 to make the color in all figure panel A show consistence. **B.** The assembly length of different assemblies after manual curation (e.g. removing duplicate regions). Numbers marked with red denote assemblies with the expected length, in the range 155,938 bp-155,945 bp. **C.** The mapping rate of validation reads to the assemblies after manual curation. Assemblies with highest mapping rate (99.43%) are marked with red. **D.** The average per-base error rate of validation reads mapped to each manually-curated genome assembly. Assemblies with the lowest error rate (0.0007) are marked with red.

**Figure S3.**
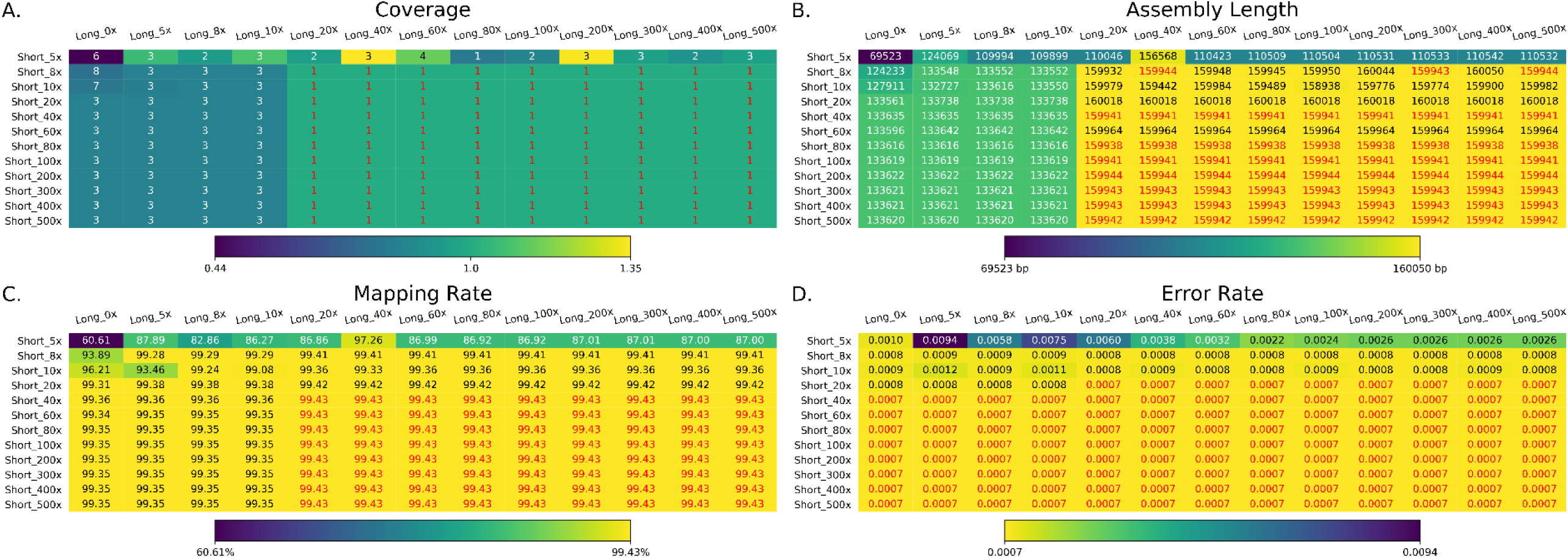
The summary of short-read-only and hybrid assemblies with SPAdes short-read correction. Long and Short indicate the coverage of this assembly. **A.** The total coverage of the chloroplast genome across all contigs output by the assembler. The number is the number of contigs of each assembly, whereas the heatmap is the genome coverage (it could be over 100% if some duplications exist). Numbers marked with red contained a single contig covering the whole chloroplast genome. The heatmap color is reversed compared to the Figure 1 to make the color in all figure panel A show consistence. **B.** The assembly length of different assemblies after manual curation (e.g. removing duplicate regions). Numbers marked with red denote assemblies with the expected length, in the range 155,938 bp-155,945 bp. **C.** The mapping rate of validation reads to the assemblies after manual curation. Assemblies with highest mapping rate (99.43%) are marked with red. **D.** The average per-base error rate of validation reads mapped to each manually-curated genome assembly. Assemblies with the lowest error rate (0.0007) are marked with red.

**Figure S4.**
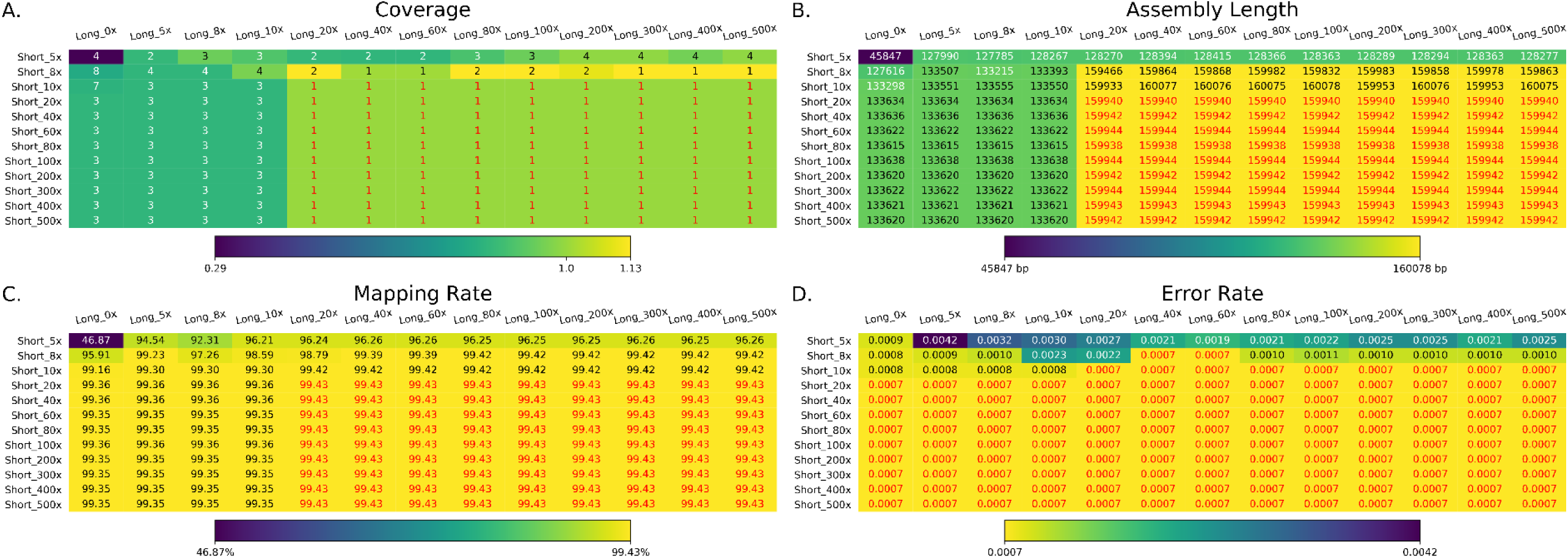
The summary of short-read-only and hybrid assemblies with Karect+SPAdes short-read correction. Long and Short indicate the coverage of this assembly. **A.** The total coverage of the chloroplast genome across all contigs output by the assembler. The number is the number of contigs of each assembly, whereas the heatmap is the genome coverage (it could be over 100% if some duplications exist). Numbers marked with red contained a single contig covering the whole chloroplast genome. The heatmap color is reversed compared to the Figure 1 to make the color in all figure panel A show consistence. **B.** The assembly length of different assemblies after manual curation (e.g. removing duplicate regions). Numbers marked with red denote assemblies with the expected length, in the range 155,938 bp-155,945 bp. **C.** The mapping rate of validation reads to the assemblies after manual curation. Assemblies with highest mapping rate (99.43%) are marked with red. **D.** The average per-base error rate of validation reads mapped to each manually-curated genome assembly. Assemblies with the lowest error rate (0.0007) are marked with red.

**Figure S5.**
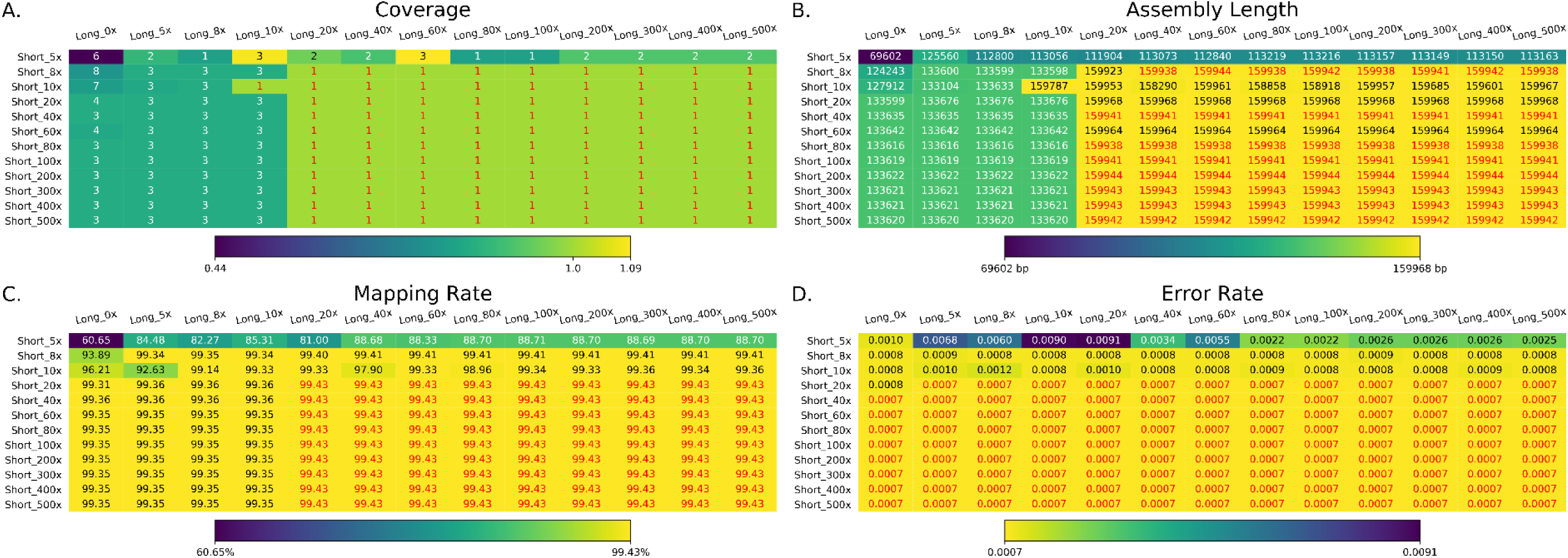
The summary of short-read-only and hybrid assemblies without any short-read correction. Long and Short indicate the coverage of this assembly. **A.** The total coverage of the chloroplast genome across all contigs output by the assembler. The number is the number of contigs of each assembly, whereas the heatmap is the genome coverage (it could be over 100% if some duplications exist). Numbers marked with red contained a single contig covering the whole chloroplast genome. The heatmap color is reversed compared to the Figure 1 to make the color in all figure panel A show consistence. **B.** The assembly length of different assemblies after manual curation (e.g. removing duplicate regions). Numbers marked with red denote assemblies with the expected length, in the range 155,938 bp-155,945 bp. **C.** The mapping rate of validation reads to the assemblies after manual curation. Assemblies with highest mapping rate (99.43%) are marked with red. **D.** The average per-base error rate of validation reads mapped to each manually-curated genome assembly. Assemblies with the lowest error rate (0.0007) are marked with red.

**Figure S6.**
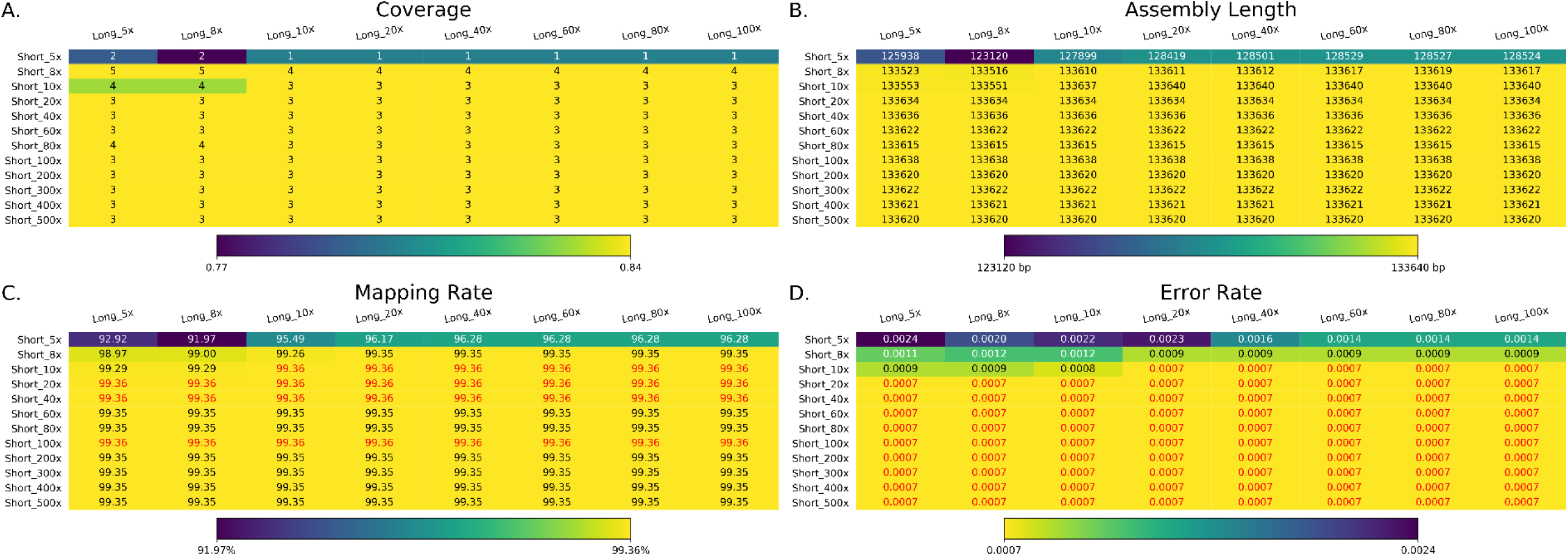
The summary of assemblies with Karect-corrected short-reads and =S10 kb long-reads. Long and Short indicate the coverage of this assembly. **A.** The total coverage of the chloroplast genome across all contigs output by the assembler. The number is the number of contigs of each assembly, whereas the heatmap is the genome coverage (it could be over 100% if some duplications exist). The heatmap color is reversed compared to the Figure 1 to make the color in all figure panel A show consistence. **B.** The assembly length of different assemblies after manual curation (e.g. removing duplicate regions). **C.** The mapping rate of validation reads to the assemblies after manual curation. Assemblies with highest mapping rate (99.43%) are marked with red. **D.** The average per-base error rate of validation reads mapped to each manually-curated genome assembly. Assemblies with the lowest error rate (0.0007) are marked with red.

**Figure S7.**
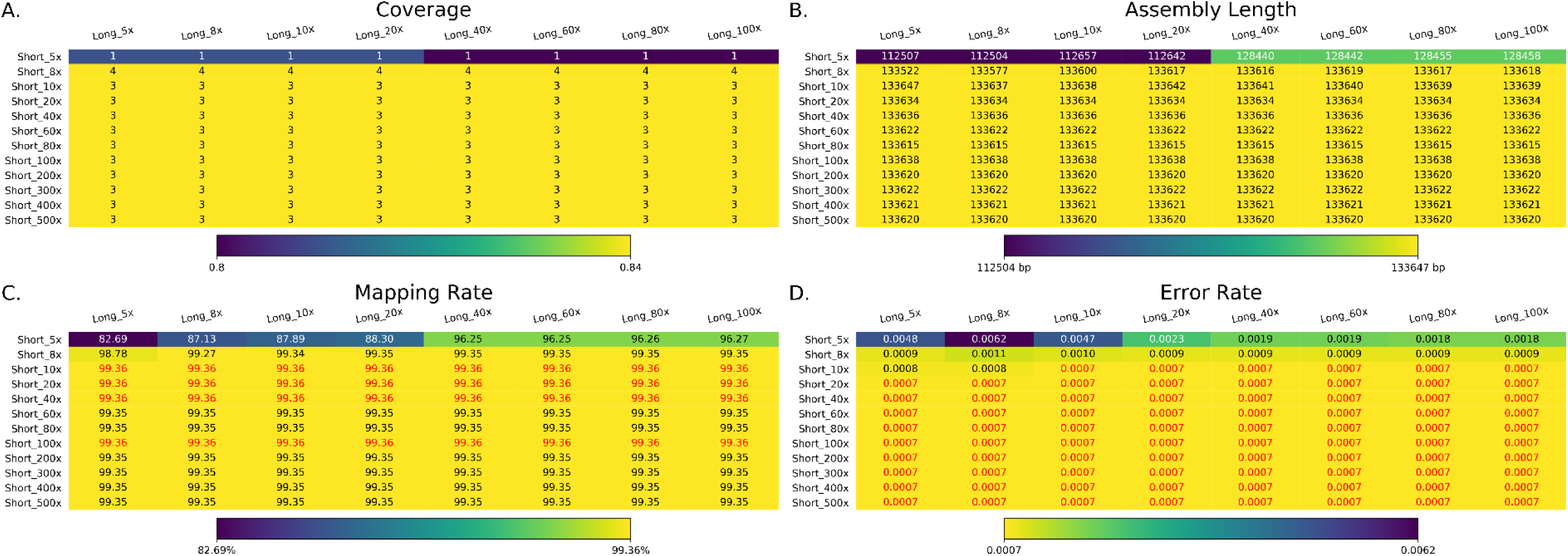
The summary of assemblies with Karect corrected short-reads and 10–20 kb long-reads. Long and Short indicate the coverage of this assembly. **A.** The total coverage of the chloroplast genome across all contigs output by the assembler. The number is the number of contigs of each assembly, whereas the heatmap is the genome coverage (it could be over 100% if some duplications exist). The heatmap color is reversed compared to the Figure 1 to make the color in all figure panel A show consistence. **B.** The assembly length of different assemblies after manual curation (e.g. removing duplicate regions). **C.** The mapping rate of validation reads to the assemblies after manual curation. Assemblies with highest mapping rate (99.43%) are marked with red. **D.** The average per-base error rate of validation reads mapped to each manually-curated genome assembly. Assemblies with the lowest error rate (0.0007) are marked with red.

**Figure S8.**
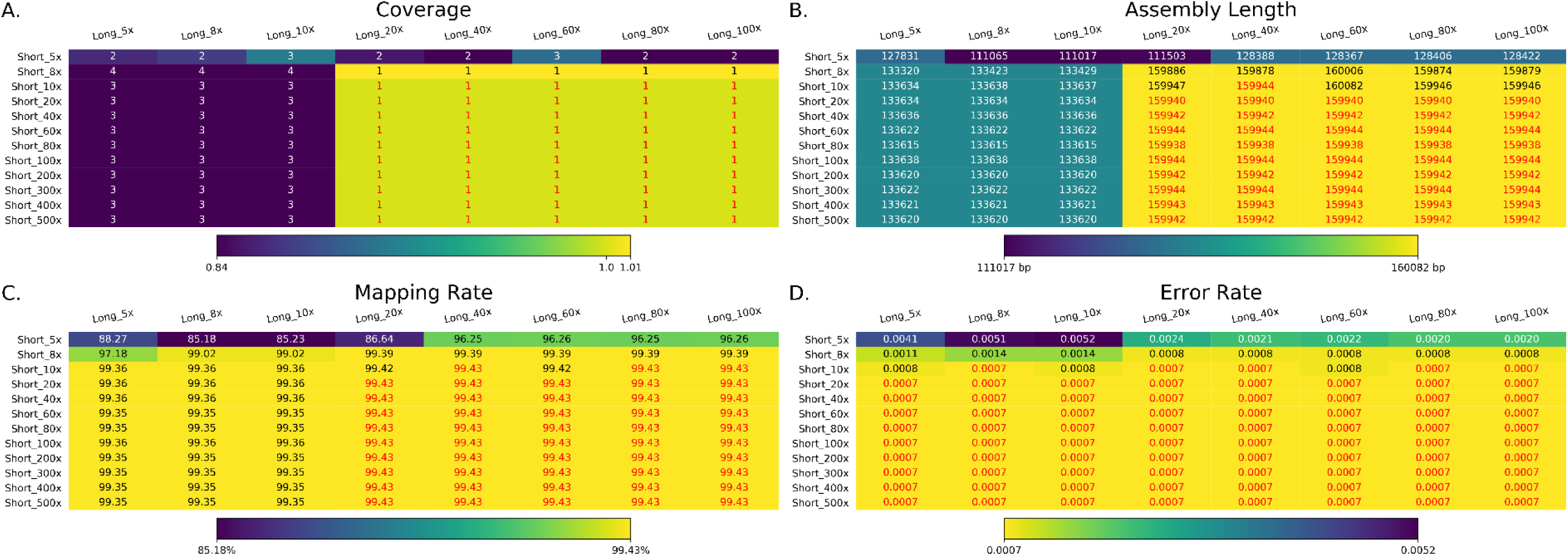
The summary of assemblies with Karect corrected short-reads and 20–30 kb long-reads. Long and Short indicate the coverage of this assembly. **A.** The total coverage of the chloroplast genome across all contigs output by the assembler. The number is the number of contigs of each assembly, whereas the heatmap is the genome coverage (it could be over 100% if some duplications exist). Numbers marked with red contained a single contig covering the whole chloroplast genome. The heatmap color is reversed compared to the Figure 1 to make the color in all figure panel A show consistence. **B.** The assembly length of different assemblies after manual curation (e.g. removing duplicate regions). Numbers marked with red denote assemblies with the expected length, in the range 155,938 bp-155,945 bp. **C.** The mapping rate of validation reads to the assemblies after manual curation. Assemblies with highest mapping rate (99.43%) are marked with red. **D.** The average per-base error rate of validation reads mapped to each manually-curated genome assembly. Assemblies with the lowest error rate (0.0007) are marked with red.

**Figure S9.**
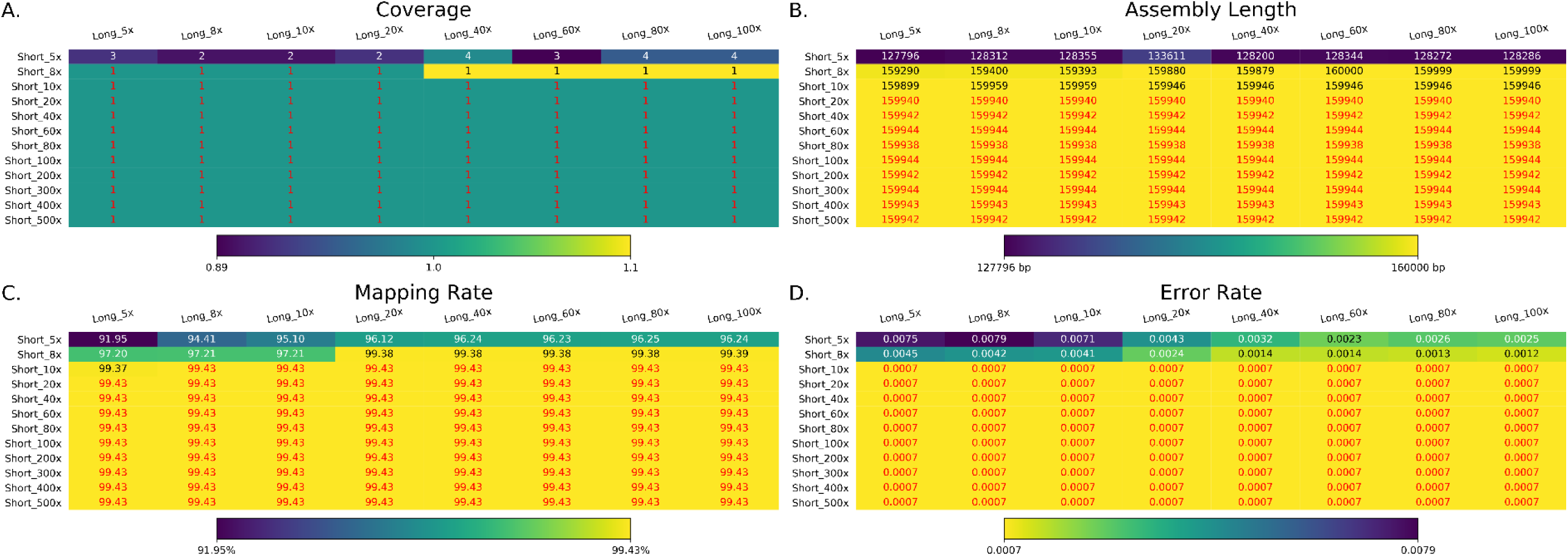
The summary of assemblies with Karect corrected short-reads and 30–40 kb long-reads. Long and Short indicate the coverage of this assembly. **A.** The total coverage of the chloroplast genome across all contigs output by the assembler. The number is the number of contigs of each assembly, whereas the heatmap is the genome coverage (it could be over 100% if some duplications exist). Numbers marked with red contained a single contig covering the whole chloroplast genome. The heatmap color is reversed compared to the Figure 1 to make the color in all figure panel A show consistence. **B.** The assembly length of different assemblies after manual curation (e.g. removing duplicate regions). Numbers marked with red denote assemblies with the expected length, in the range 155,938 bp-155,945 bp. **C.** The mapping rate of validation reads to the assemblies after manual curation. Assemblies with highest mapping rate (99.43%) are marked with red. **D.** The average per-base error rate of validation reads mapped to each manually-curated genome assembly. Assemblies with the lowest error rate (0.0007) are marked with red.

**Figure S10.**
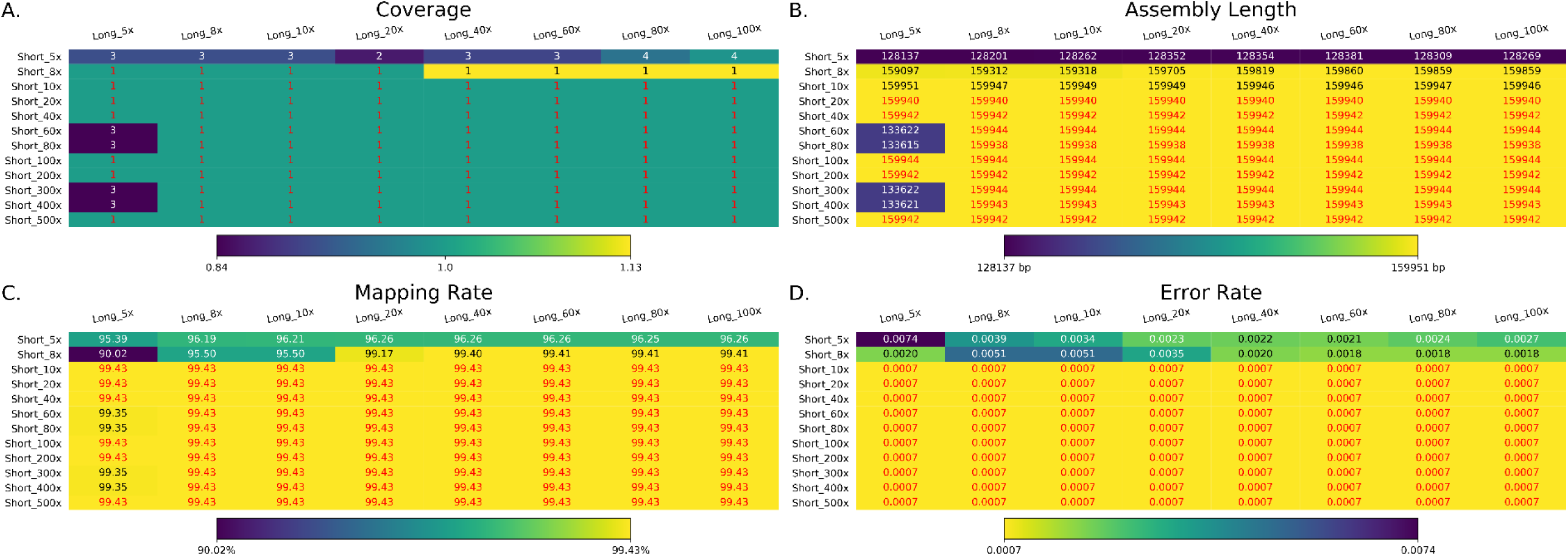
The summary of assemblies with Karect corrected short-reads and 40–50 kb long-reads. Long and Short indicate the coverage of this assembly. **A.** The total coverage of the chloroplast genome across all contigs output by the assembler. The number is the number of contigs of each assembly, whereas the heatmap is the genome coverage (it could be over 100% if some duplications exist). Numbers marked with red contained a single contig covering the whole chloroplast genome. The heatmap color is reversed compared to the Figure 1 to make the color in all figure panel A show consistence. **B.** The assembly length of different assemblies after manual curation (e.g. removing duplicate regions). Numbers marked with red denote assemblies with the expected length, in the range 155,938 bp-155,945 bp. **C.** The mapping rate of validation reads to the assemblies after manual curation. Assemblies with highest mapping rate (99.43%) are marked with red. **D.** The average per-base error rate of validation reads mapped to each manually-curated genome assembly. Assemblies with the lowest error rate (0.0007) are marked with red.

**Figure S11.**
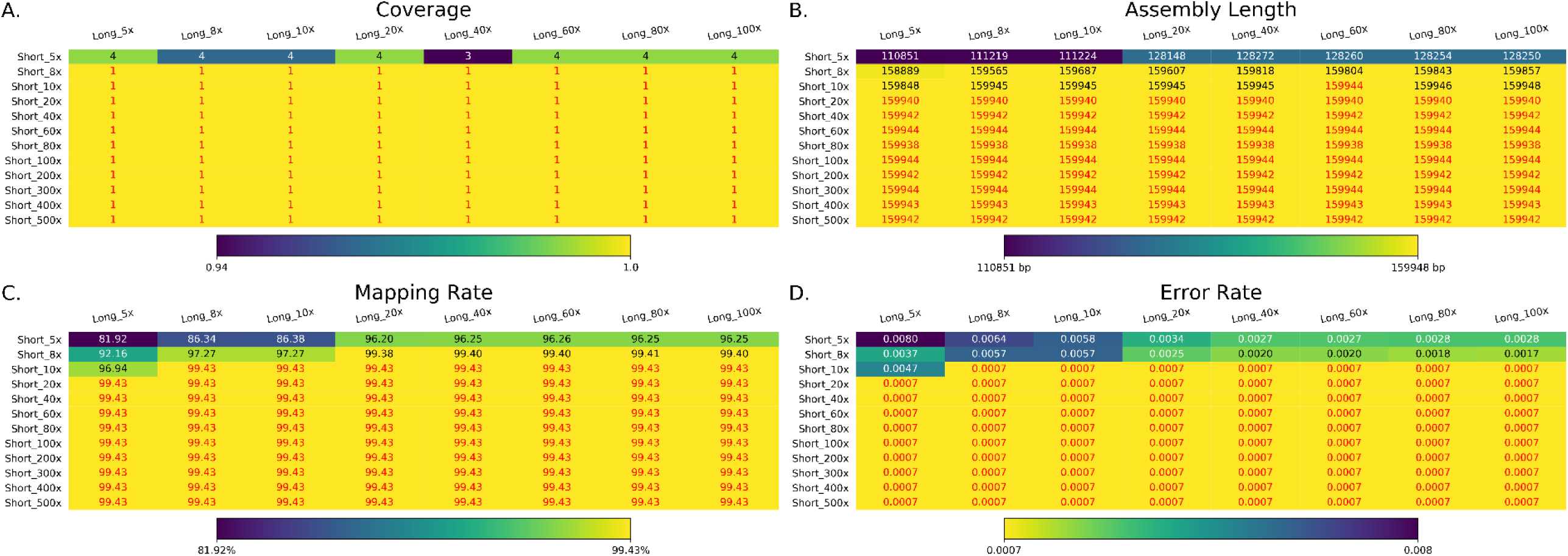
The summary of assemblies with Karect corrected short-reads and 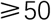 kb long-reads. Long and Short indicate the coverage of this assembly. **A.** The total coverage of the chloroplast genome across all contigs output by the assembler. The number is the number of contigs of each assembly, whereas the heatmap is the genome coverage (it could be over 100% if some duplications exist). Numbers marked with red contained a single contig covering the whole chloroplast genome. The heatmap color is reversed compared to the Figure 1 to make the color in all figure panel A show consistence. **B.** The assembly length of different assemblies after manual curation (e.g. removing duplicate regions). Numbers marked with red denote assemblies with the expected length, in the range 155,938 bp-155,945 bp. **C.** The mapping rate of validation reads to the assemblies after manual curation. Assemblies with highest mapping rate (99.43%) are marked with red. **D.** The average per-base error rate of validation reads mapped to each manually-curated genome assembly. Assemblies with the lowest error rate (0.0007) are marked with red.

**Figure S12.**
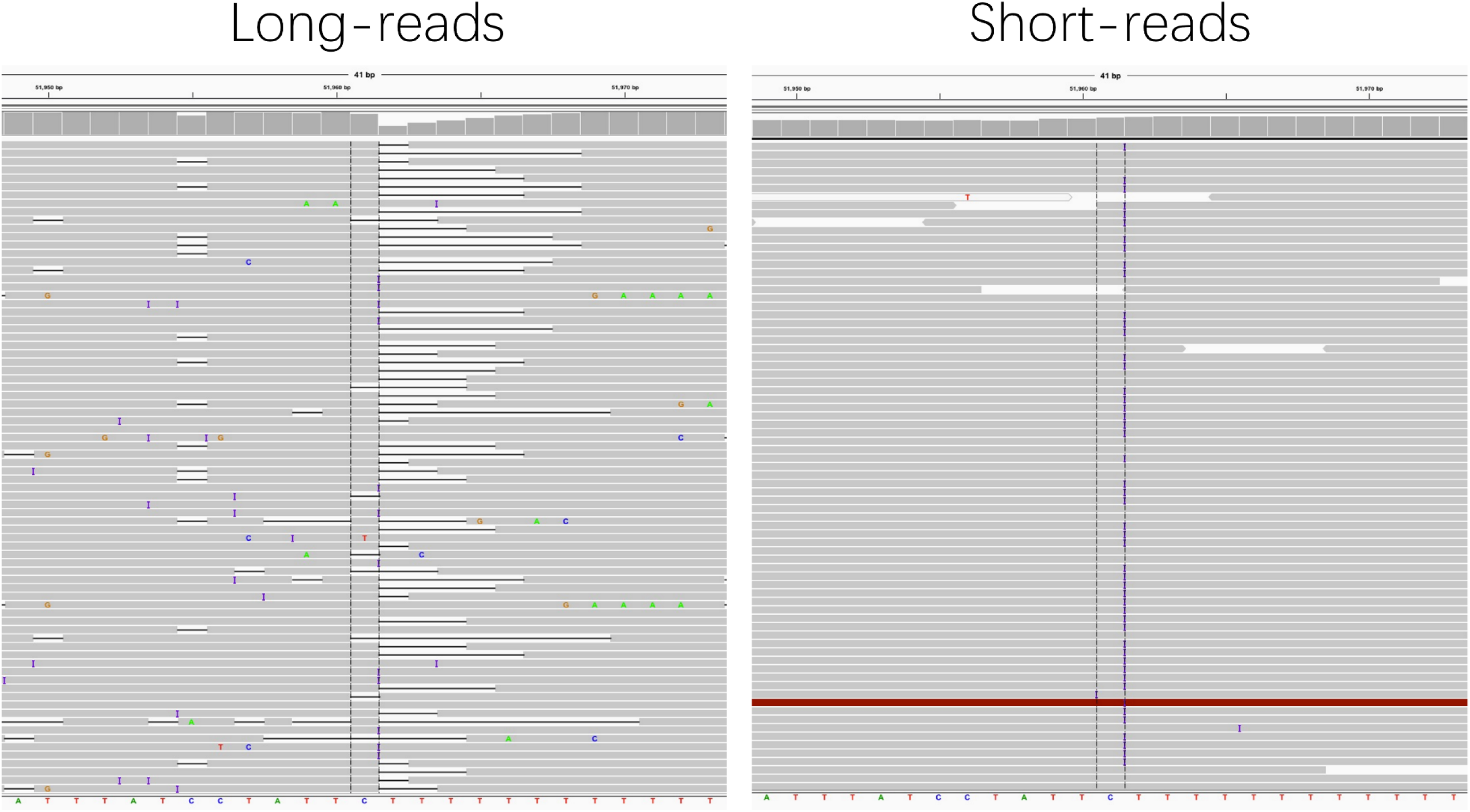
The long-/short-read mapping coverage in the possible heteroplastic site 51961 (IGV view). The purple "I” in short-reads indicated the T insertion, whereas the black lines in long-reads indicate the deletion during that region.

